# Ancestry Composition: A Novel, Efficient Pipeline for Ancestry Deconvolution

**DOI:** 10.1101/010512

**Authors:** Eric Y. Durand, Chuong B. Do, Joanna L. Mountain, J. Michael Macpherson

**Affiliations:** 23andMe, Inc., Mountain View, CA, USA; School of Computational Sciences, Chapman University, Orange, CA, USA

## Abstract

Ancestry deconvolution, the task of identifying the ancestral origin of chromosomal segments in admixed individuals, has important implications, from mapping disease genes to identifying candidate loci under natural selection. To date, however, most existing methods for ancestry deconvolution are typically limited to two or three ancestral populations, and cannot resolve contributions from populations related at a sub-continental scale.

We describe Ancestry Composition, a modular three-stage pipeline that efficiently and accurately identifies the ancestral origin of chromosomal segments in admixed individuals. It assumes the genotype data have been phased. In the first stage, a support vector machine classifier assigns tentative ancestry labels to short local phased genomic regions. In the second stage, an autoregressive pair hidden Markov model simultaneously corrects phasing errors and produces reconciled local ancestry estimates and confidence scores based on the tentative ancestry labels. In the third stage, confidence estimates are recalibrated using isotonic regression.

We compiled a reference panel of almost 10,000 individuals of homogeneous ancestry, derived from a combination of several publicly available datasets and over 8,000 individuals reporting four grandparents with the same country-of-origin from the member database of the personal genetics company, 23andMe, Inc., and excluding outliers identified through principal components analysis (PCA). In cross-validation experiments, Ancestry Composition achieves high precision and recall for labeling chromosomal segments across over 25 different populations worldwide.

## 1 Introduction

The genome of an admixed individual can be viewed as a mosaic of chromosomal segments of different ancestry (Falush *et al.*, 2003; Tang *et al.*, 2006). Ancestry deconvolution, or *local ancestry inference*, is the task of resolving the ancestral origin of such segments. Robust ancestry deconvolution fundamentally enables several important lines of research, including admixture-based mapping of diseases (Seldin *et al.*, 2011), controlling for population structure in disease-association studies (Price *et al.*, 2010), and studies of population history (Novembre and Ramachandran, 2011; Hellenthal *et al.*, 2014).

A number of methods that successfully perform ancestry deconvolution have been introduced; perhaps the most widely-used are `structure`, LAMP-ANC/LAMP-LD, and `HAPMIX` (Falush *et al.*, 2003; Pasaniuc *et al.*, 2009; Price *et al.*, 2009). The setting in which these methods achieve their success has been limited to resolving ancestry from a small number of populations separated by comparatively large genetic, and therefore typically geographic, distances. This we refer to as *continental-scale* separation, e.g., the respective distances separating European, West African and Amerindian populations. None of the methods have been shown to be successful in resolving ancestry from many subcontinental-scale populations, e.g., Northern European vs. Southern European, in an individual at once. `HAPMIX` and LAMP-ANC/LAMP-LD do not allow more than 2 and 5 source populations, respectively, to be considered simultaneously. Of the three methods, only LAMP-ANC/LAMP-LD has been shown to be effective at resolving subcontinental-scale ancestry, although this was in a setting restricted to pairwise comparisons (Pasaniuc *et al.*, 2009).

Two recent methods have made progress in subcontinental-scale ancestry deconvolution (Omberg *et al.*, 2012; Maples *et al.*, 2013). These methods introduced classification techniques from machine learning, namely support vector machines (Omberg *et al.*, 2012) and random forests (Maples *et al.*, 2013). Both methods demonstrate some ability to resolve subcontinental-scale ancestry with simulated admixtures of pairs of populations. But no method has yet shown the ability to resolve contributions from several subcontinental-scale source populations at once.

We are concerned here with deconvolving ancestry from multiple human source populations separated by subcontinental-scale genetic/geographic distances, with particular attention to European-Americans. Any practical method for resolving ancestry at such distances must be capable of considering many possible source populations simultaneously. While it is relatively uncommon for an individual to have ancestors from more than two or three populations separated by continental-scale distances, it does not appear to be uncommon for individuals to have genetic contributions from multiple populations separated by subcontinental-scale distances (Seldin *et al.*, 2006; Price *et al.*, 2008; Nelis *et al.*, 2009). Further, in this context, it may be difficult to know *a priori* which subcontinental source populations have contributed to a given individual’s genome, so it is advantageous to be able to consider many possible sources simultaneously.

In this article, we introduce a new ancestry deconvolution method, *Ancestry Composition*. Ancestry Composition is a modular three-stage pipeline. First, preliminary ancestry assignments are obtained via support vector machines using a string kernel. Next, these preliminary assignments are processed using a novel autoregressive pair hidden Markov model that corrects misassignments and phasing errors. Last, it employs isotonic regression to calibrate the assignments globally, which controls overall false positive and false negative error rates. This is the first time a recalibration method has been used in ancestry inference. Ancestry Composition achieves high precision and recall for labelling chromosomal segments from over 25 populations worldwide. We show also that Ancestry Composition can successfully resolve ancestry from multiple subcontinental European populations simultaneously.

## 2 Methods

Ancestry Composition is composed of three largely independent modules:

1. **Local classifier module:** A kernel-based support vector machine classifier for assigning tentative ancestry labels to short local phased genomic regions.
2. **Error correction module:** An autoregressive pair hidden Markov model (pair-HMM) that takes the tentative ancestry labels from Step 1 as input, and produces reconciled local ancestry estimates with assigned confidence scores, while simultaneously correcting phasing errors.
3. **Recalibration module:** An isotonic regression procedure for recalibrating the confidence estimates from Step 2.

The overall pipeline is illustrated in Figure 1. We describe each of these components in the following subsections.

**Figure 1:**
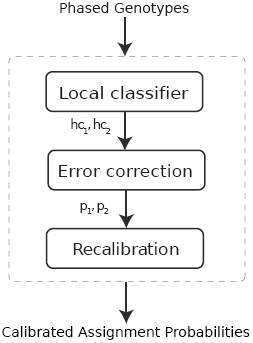
Overview of Ancestry Composition pipeline. An individual genotype is first phased using an out-of-sample extension of BEAGLE, which outputs two haplotypes, *hap*_1_*, hap*_2_. The haplotypes are passed to the local classifier, where they are split in equal size chunks that are classified independently to one of the reference populations. The local classifier outputs two hard-clustering vectors, *hc*_1_*, hc*_2_, composed of populations of origin on each haplotype. The error correction module takes *hc*_1_*, hc*_2_ as input, reconciles hard-clustering calls and corrects for switch errors. It outputs two vectors, *p*_1_*, p*_2_, containing assignment probabilities along both haplotypes. Finally, *p*_1_ and *p*_2_ are fed to a recalibration module that outputs assignment confidence scores along both haplotypes.

### 2.1 Local classifier module

The task of the local classifier is to assign each marker along both haplotypes to one of *K* reference populations. The local classifier starts by splitting each haplotype into *S* segments of *M* biallelic markers (*M* was set to 100 in our experiments). Each segment is treated independently and is assumed to have a single ancestral origin. Thus, the local classifier returns a vector (*hc*_*i*_)_1:*S*_ where *hc*_*i*_ ∈ {1 *… K*} is the hard-clustering value assigned to segment *i*. We implemented the local classifier using a support vector machine (SVM) classifier using a specific choice of string kernel, as described below. Note that the local classifier module requires haplotypes to be pre-phased. Phasing uncertainty is not taken into account in the local classifier, but is dealt with in the error correction module.

#### 2.1.1 Kernel support vector machines

Support Vector Machines (SVMs) are a class of supervised learning algorithms first introduced by (Vapnik, 1998). In their most basic form, SVMs are non-probabilistic binary linear classifiers, i.e., they learn a linear decision boundary that can be used to discriminate between two classes. SVMs can be extended to problems that are not separable by a hyperplane, using the soft-margin technique (Cristianini and Shawe-Taylor, 2000).

Consider a set of training data 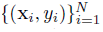 where x_*i*_ is some feature vector in ℝ*^d^*, *y*_*i*_ ∈ {0, 1} its corresponding class, *d* is the dimension of the feature vectors and *N* is the number of training chromosomes. The SVM learns the decision boundary by solving the quadratic programming optimization problem (1):

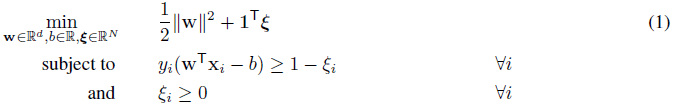

**Encoding the feature vectors.** In our application, each feature vector x_*i*_ represents the encoding of a segment of *M* biallelic markers from a pre-phased haplotype. One natural encoding is to use one feature per marker, each feature then encoding the presence/absence of the minor allele. However, this encoding fails to capture the spatial relationship of consecutive markers within the segment, *i.e.* the linkage pattern, a distinguishing feature of haplotypes. Instead, we use every possible *k*-mer (*k* ∈ {1 *… M*}) as our features. For a segment of *M* biallelic markers, there are *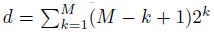* possible *k*-mers, which implies on the order of 10^30^ features when *M* = 100. It is not feasible to directly construct feature vectors with this many dimensions in memory; in the next section, however, we introduce a string kernel in order to facilitate working with our high-dimensional feature set.

**String kernel.** A key feature of SVMs is that solving (1) is equivalent to solving the dual quadratic programming problem (2):

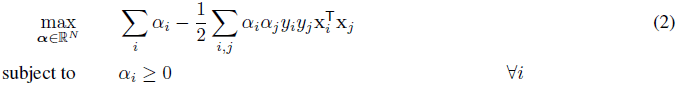

The dual representation of the SVM optimization problem is only dependent on the inner product 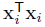, and thus allows for the introduction of kernels (Boser *et al.*, 1992). Kernels offer a way of mapping observations to a high dimension feature space, and can allow for a huge computational advantage as they can be evaluated without explicitly calculating feature vectors. Let us denote *χ* the input space and *ϕ*: *χ →* {0, 1}*^d^* the mapping such that for any segment *x* of length *M*, *ϕ*(*x*) is the vector whose elements denote the presence/absence of each of the *d* possible *k*-mers in *x*. We then define our string kernel as, Ɐ*i, j* ∈ {1*, …, N*},

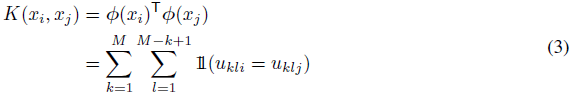

where *u*_*kli*_ is the *k*-mer starting at position *l* in haplotype segment *i*. Our kernel is a special case of the weighted degree kernel (Rätsch *et al.*, 2006). Standard dynamic programming techniques can be used to evaluate *K*(*x*_*i*_, *x*_*j*_) in *O*(*M*) operations without explicitly enumerating all *d* features for each mapped input vector. Thus, the string kernel allows extraction of a large amount of information from the haplotype segments without compromising the speed of the local classifier and without explicitly computing the feature vectors.

**Multiclass SVM.** SVMs are fundamentally binary classifiers, but in this setting we are concerned with deciding not between two but between 25 possible populations. To assign a single hard-clustering value *k* ∈ {1*, …, K*} to a haplotype segment, we trained 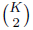 classifiers, one for each pair of populations. We then assigned the haplotype to a single population using a straight-forward majority vote across all pairs. We also experimented with a one-vs-all approach, which did not perform as well as the one-vs-one approach we took.

#### 2.1.2 Training data

We trained the local classifier on more than 9700 individuals divided into *K* = 25 reference populations. We denote *N* the number of training individuals. The reference populations are a combination of countries and broader geographic regions. We required that a population contain at least 25 individuals. When a country did not contain enough individuals, it was grouped with countries from the same geographic region. The process was repeated until each population contained enough individuals. Next, we describe the steps involved in building the training set.

We started by building a dataset of *N* ^*^ = 10699 unrelated individuals with known/self-reported ancestry. Out of the 10699 individuals, 1793 came from publicly available datasets (1000 Genomes: 765, CEPH-HGDP: 941, HapMap3: 87).

The remaining 8906 were research-consented 23andMe members who reported via a survey on the 23andMe website that their four grandparents were born in the same country. The text of the questions asked was “In which country was your (mother’s mother) born?", for each of the four grandparents. The answer was chosen from a list of countries, which included the option “I don’t know”. This number includes 23andMe members who indicated that all four of their grandparents had Ashkenazi Jewish ancestry. This was ascertained by a separate question on the same survey. The text of that question reads: “If your (mother’s mother’s) birthplace does not fully describe (her) ancestry, please provide additional ancestry-relevant information here. Your answer may be any mix of national, ethnic, religious, or other labels, separated by semicolons. To illustrate: transnational or ethnic identities, such as *Ashkenazi*, *Basque*, and *Cherokee*, are often more relevant to ancestry than birthplace. Or, birthplace may not accurately reflect ancestry for a child of immigrants. For example, if this person was born in the US to immigrants from India, you might write *Hindu*; *Indian*.” The text supplied by members in response to this question was searched for occurrences of “Ashkenazi".

We ensured that all the reference individuals were distantly related by removing individuals from the sample until no two individuals shared more than an estimated 100 cM identical-by-descent (IBD). IBD estimates were obtained according to Henn *et al.* (2012).

We then performed genome-wide PCA on the remaining individuals. We computed the distance of each individual to his/her 20 nearest-neighbors in the space defined by the first ten principal components (PCs), which enabled us to infer the empirical distribution of distance to nearest neighbors for each of the *K* populations. Finally, we removed individuals outside of the 5th-quantile interval of the distribution. Table S1 and Table S2 summarize the composition of our training set.

### 2.2 Error correction module

The local classifier provides noisy estimates of a segment’s ancestry. The role of error correction is to smooth the local classifier’s estimates, using information from adjacent segments on the same chromosome and on the opposing chromosome. It uses hard clustering assignments from the SVM step as input and outputs assignment probabilities for each segment. We have implemented the ancestry correction module as an *autoregressive pair hidden Markov model* (APHMM). Recall that *S* denotes the number of segments of *M* markers, and consider a directed probabilistic graphical model consisting of:

- *S* hidden states (*y*_1:*S*_ = (*y*_1_*, y*_2_*, …, y_S_*)), where 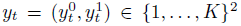 represents the true population labels of the pair of haplotype segments at a particular position *t* in the genome,
- *S* observed states (*x*_1:*S*_ = (*x*_1_*, x*_2_*, …, x_S_*)), where 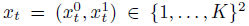 represents the observed population labels for the pair of haplotype segments at position *t*, i.e., the output from the local classifier.
- *S* − 1 hidden switch indicators (*s*_2:*S*_ = (*s*_2_*, s*_3_*, …, s_S_*)), where *s*_*t*_ ∈ {0, 1} denotes whether a switch error has occurred between *x*_*t*-1_ and *x*_*t*_.

In this model, we explicitly represent the segments corresponding to both haplotypes covering a genomic region. We implicitly assume that switch errors in the initial phasing occur only at the boundaries of segments, and we model the joint probability of *y*_1:*S*_, *x*_1:*S*_, and *s*_2:*S*_ as

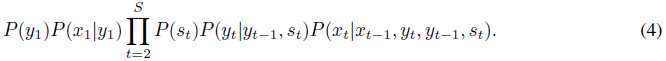

Figure S1 illustrates the graphical model for a sequence of length *S* = 5.

#### 2.2.1 Model parameterization

The parameters of our model are:

- *μ*_1_*, …, μ_K_*: the prior distribution for hidden states,
- *μ*_1|*y*_*, …, μ*_*K*|*y*_: the prior distribution for emissions conditional on hidden state *y*,
- *σ*: the prior probability that a switch error occurs between any two given consecutive positions,
- *θ*: the prior probability of a recombination between two consecutive positions
- *∈_y,x_*: the prior probability of a label reset between two consecutive positions with the same hidden state *y*, where the former label is *x*

In total, there are 2*K*^2^ + 1 free parameters in our model.

Below, we define each of the components of the joint probability expression (4) in terms of the free parameters of the model.

1. **Initial state distribution.** We assume that the population assignments for each haplotype are sampled independently from the set of global admixture proportions:

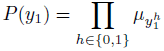
2. **Initial emission distribution.** Like the initial state, the initial emissions for each haplotype are sampled independently from a set of global label probabilities *μ*_1|*i*_, …, *μ*_*K*|*i*_ conditional on the underlying true population *i*:

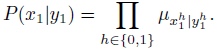
3. **Switch error model.** We assume that switch errors occur with constant probability *σ* between each pair of states:

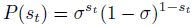
4. **Transition probability model.** In the transition probability model, we consider each haplotype separately. In each haplotype, a recombination occurs with probability *?*, which results in a new hidden population label being drawn from *μ*_1_*, …, μ_K_*.

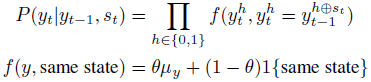
5. **Emission probability model.** In the emission probability model, we consider each haplotype independently. In each haplotype, if the current hidden population label is the same as the previous hidden population label, then a reset occurs with probability *o*, which results in a new observed population label being drawn from 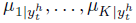. However, if the current hidden population label has changed from the previous hidden population label, then a new observed population label is drawn from *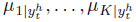*.

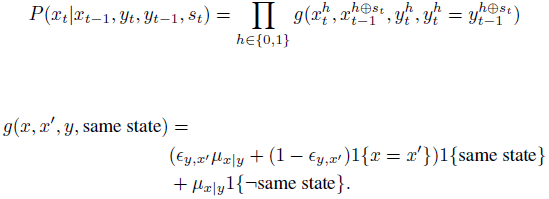

The parameters of the model can be estimated using the standard expectation maximization (EM) algorithm (Dempster *et al.*, 1977). Posterior probabilities for each segment are estimated using the well-known forward and backward algorithms for hidden Markov models. Using dynamic programming techniques, the complexity of the posterior decoding step is *O*(*LK*^2^), where *L* is the number of windows to decode and *K* is the number of populations.

### 2.3 Recalibration module

To explore the degree to which the assignment probabilities emerging from the error correction module were well-calibrated, we employed so-called reliability plots. *Reliability* is defined as the degree of correspondence between assignment probabilities and observed assignment accuracy (Murphy and Winkler, 1977). In a reliability plot, assignment probabilities are plotted against empirical class membership probabilities (Niculescu-mizil and Caruana, 2005); departure from a straight line from (0, 0) to (1, 1) is a visual indication of miscalibration.

Finding evidence of miscalibration within some populations, we employed isotonic regression to recalibrate the ancestry correction module assignment probabilities. Isotonic regression models are a particular type of one-dimensional regression model in which the response variable is a modeled as an arbitrary non-decreasing function of a single input feature (Zadrozny and Elkan, 2002). We implemented a regularized version of isotonic regression in which we solve the following quadratic programming problem:

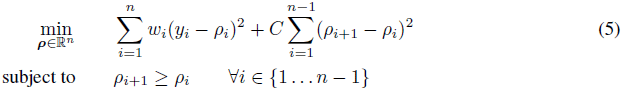

where *y*_*i*_ are the probabilities to be recalibrated (i.e., the output of the error correction module), *w*_*i*_ are weights and *C* is a penalty parameter (set to 0.001 in our experiments). We solved (5) using the `CVXOPT` package (Andersen *et al.*, 2011), thus estimating *ρ_s_* = (*ρ*_*s*1_, …, *ρ*_*sK*_) for each segment *s*. Finally, we renormalized *ρ_s_* so that 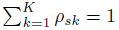 The output of the recalibration module consists in recalibrated posterior probabilities for each segment *s* ∈ {1*, …, S*}.

**Hierarchical classifier.** Populations from the same geographical area are not always easy to distinguish. For instance, it might not always be possible to confidently decide whether a segment originated from Scandinavia or the British Isles, either because we lack power due to small reference populations, or because the corresponding haplotypes are truly at similar frequencies in the two populations. In such a case, however, it may be possible to decide that that segment originated from North Europe rather than South Europe, even if the finer distinction cannot be made. To address this issue, we define a population hierarchy that groups population from the same area together (Figure S2). The resulting hierarchy has four levels. The *K* leaves of the hierarchy correspond to the *K* reference populations. The top level consists of a single root node representing the union of all populations. Broadly, the levels beneath that correspond to continent-scale, regional-scale, and sub-regional–scale distinctions. Leaf/terminal nodes may occur at any of these levels; for example, South Asia and Oceania are in the first level of the hierarchy, so at the continent scale, but are not further subdivided.

In order to estimate assignment probabilities to internal nodes for a segment *s*, we sum the recalibrated posteriors *ρ_s_* from the *K* leaves to the root of the tree, so that each tree node is associated with a probability equal to the sum of its children’s probabilities. The probability at the root is always 1 because we renormalized *ρ_s_*. The recalibration procedure ensures that the precision of predicting that a segment originated from node *o* is at least *t* if 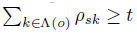, where Λ(*o*) is the set of leaves descending from node *o*.

Therefore, we can choose a desired precision level, for example *t* = 0.8. We then sum the recalibrated posterior probabilities up the tree, from the *K* = 25 leaves to the root. We then assign the segment to the lowest (i.e., closest to the leaves) node at which the posterior probability exceeds *t*. In situations where more than one node fits the above criteria, we choose the qualifiying node with the highest posterior probability. In the worst case, no node other than the root satisfies the criterion, in which case the segment is unclassified. When assignment probabilities are well calibrated, this procedure ensures that the precision of the assignment is at least *t*. Therefore, we refer to *t* as the precision threshold in what follows.

### 2.4 Model estimation and evaluation

For model estimation and evaluation, we used a multi-stage cross-validation procedure, with stages corresponding to the three modules within the Ancestry Composition pipeline.

For the local classifier module, we employed a stratified cross-validation procedure in which we split the reference individuals into five equally-sized groups (while maintaining similar representation among the *K* = 25 reference populations within each group). For each of these groups, we evaluated the local classifier module using model parameters learned using data from the other four groups.

For the error correction module, we again used a five-fold cross-validation procedure when estimating emission parameters for the autoregressive pair-HMM. Within each fold, we estimated transition parameters by an unsupervised EM training procedure that relied on the natural admixture found in a broader set of admixed 23andMe customers. To allow for heterogeneity in transition parameters by general ancestry, we clustered general 23andMe customers into a small number of population groups based on principal components analysis (PCA), fit separate transition parameters in each population group, and then combined the predictions from each separation model using Bayesian model averaging. Specifically, we fit transition parameters using 1,000 unrelated 23andMe customers from each of six different population groups: Latinos, African Americans, Europeans, Ashkenazi Jewish, East Asians and South Asians.

Finally, for the recalibration model, we fit isotonic regression models using a combination of data from individuals of homogeneous ancestry from the reference populations and individuals of heterogeneous ancestry using a forward-in-time admixture simulation, drawing segments from the reference individuals, assuming a uniform recombination rate of *ρ* = 10^-8^ per base pair and 10 generations since admixture. We note that the goal of this procedure is not to produce realistic simulations, but rather to fit isotonic regression models on individuals with many different ancestry transition backgrounds.

## 3 Implementation

### 3.1 Effects of recalibration module

We applied our recalibration procedure to each of the 25 reference populations. To measure the degree of miscalibration before and after applying our recalibration procedure, we calculated the recalibration “net difference”, defined as 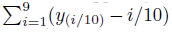, where *y*_(*i/*10)_ is the population-average assignment probability at true ancestry proportion *i/*10. There is considerable variety in the degree of miscalibration: some populations are already fairly well-calibrated leaving the error correction module, but others are miscalibrated (Table S3). When miscalibrated, the error correction module is over-confident, corresponding to a negative net difference, about as often as it is under-confident, corresponding to a positive net difference. Illustrative examples of both forms of miscalibration are provided in Fig. S3.

As may be seen in Table S3, after application of the recalibration module, none of the estimates remain over-confident, apart from what appears to be very slight over-confidence for the Mongolia population. Most of the under-confident populations become better-calibrated, in that the magnitude of their net difference is reduced, but several remain under-confident.

The preponderance of near-zero post-calibration net difference values in Table S3 suggest that our recalibration procedure indeed improves calibration overall. That several populations remain under-confident is undesirable, but this is preferable to having over-confident predictions. This under-confidence makes our accuracy measures at a given precision threshold somewhat conservative, as may be seen below.

### 3.2 Ancestry prediction performance

We evaluated Ancestry Composition’s performance as a classifier in terms of precision and recall via a fivefold stratified cross-validation experiment (see Methods). Precision for population *k* was computed as the proportion of segments predicted to be from *k* that actually are from *k*. Recall for population *k* was computed as the proportion of segments truly from population *k* that are predicted to be from population *k*.

We present the accuracy results at three different spatial scales, from coarsest-to finest-resolution: *continental*, *regional*, and *sub-regional* scales. The population hierarchy described above was used to map from the assignments, which are fundamentally made at the leaf nodes in the hierarchy, to their parent groups.

In each of these sets of results, precision and recall are shown for two precision thresholds *t*, namely *t* = 0 and *t* = 0.8; any *t* ∈ [0, 1] may be used. A segment will receive an assignment only if its posterior probability for some population meets or exceeds the precision threshold. With *t* = 0, this means that all segments will receive an assignment, even though the largest posterior probability may be very small. Informally, we term this “best-guess” mode, to capture that the classifier is effectively forced to make an assignment. With *t* = 0.8, segments may go unassigned. We note that the reported precision values cannot be considered independently from the precision threshold. Specifically, because the posterior probabilities have been calibrated, the estimated precision for some choice of *t* will never be less than *t*. The estimated precision can, however, be greater than *t*; this would simply reflect a population for which the posterior segment probabilities often exceed *t*. Recall also depends on the choice of *t*, but less directly. Generally, raising the precision threshold increases precision at the expense the recall.

At the continental scale, Ancestry Composition achieves precision of 98% or above, and recall exceeding 94%, respectively, for all but the “Middle East and Northern Africa” population (Table 1). For Europe, East Asia and America, and Sub-Saharan Africa, precision and recall are near to 99%. These values are averages over the five cross-validation folds. At this scale, performance is fairly insensitive to the choice of precision threshold *t*. Increasing *t* from 0 to 0.8 pushes precision up very slightly, and recall down, apart from Middle East and Northern Africa, where recall falls to 76%.

**Table 1:** Ancestry Composition accuracy at all scales. Precision and recall are shown for the 31 populations, evaluated at precision thresholds of *t* = 0 and *t* = 0.8. A threshold of *t* = 0 corresponds to making best-guess predictions. The values in parentheses are the respective standard errors computed across chromosomes. The “—” symbol indicates that precision and recall values are integrated over all children of the parent population.

| Continent | Region | Sub-Region | $t = 0$ | | $t = 0.8$ | |
| --- | --- | --- | --- | --- | --- | --- |
|  |  |  | Precision | Recall | Precision | Recall |
| Oceania | — | — | 0.994 (0.020) | 0.961 (0.007) | 0.999 (0.004) | 0.952 (0.011) |
| East Asia and Americas | — | — | 0.990 (0.003) | 0.993 (0.005) | 0.991 (0.003) | 0.987 (0.005) |
|  | Americas | — | 0.989 (0.004) | 0.935 (0.019) | 0.990 (0.004) | 0.858 (0.026) |
|  | Northeast Asia | — | 0.956 (0.006) | 0.989 (0.003) | 0.968 (0.004) | 0.889 (0.023) |
|  |  | Japan | 0.967 (0.025) | 0.953 (0.016) | 0.979 (0.014) | 0.917 (0.042) |
|  |  | China | 0.916 (0.014) | 0.963 (0.009) | 0.934 (0.009) | 0.913 (0.024) |
|  |  | Yakut | 0.870 (0.099) | 0.862 (0.076) | 0.958 (0.049) | 0.779 (0.107) |
|  |  | Mongolia | 0.812 (0.042) | 0.647 (0.102) | 0.890 (0.034) | 0.526 (0.095) |
|  |  | Korea | 0.763 (0.046) | 0.857 (0.103) | 0.860 (0.034) | 0.617 (0.016) |
|  | Southeast Asia | — | 0.921 (0.020) | 0.748 (0.018) | 0.950 (0.015) | 0.695 (0.028) |
| Europe | — | — | 0.989 (0.003) | 0.990 (0.002) | 0.990 (0.000) | 0.986 (0.004) |
|  | Ashkenazi | — | 0.970 (0.003) | 0.966 (0.011) | 0.972 (0.004) | 0.933 (0.018) |
|  | Northern Europe | — | 0.905 (0.022) | 0.946 (0.012) | 0.950 (0.007) | 0.848 (0.027) |
|  |  | Finland | 0.933 (0.015) | 0.908 (0.018) | 0.953 (0.010) | 0.858 (0.025) |
|  |  | Scandinavia | 0.715 (0.045) | 0.636 (0.072) | 0.858 (0.026) | 0.345 (0.067) |
|  |  | Britain/Ireland | 0.704 (0.042) | 0.828 (0.028) | 0.902 (0.021) | 0.395 (0.070) |
|  |  | France/Germany | 0.640 (0.039) | 0.390 (0.086) | 0.781 (0.03) | 0.083 (0.017) |
|  | Southern Europe | — | 0.851 (0.025) | 0.842 (0.047) | 0.928 (0.008) | 0.660 (0.073) |
|  |  | Sardinia | 0.859 (0.089) | 0.810 (0.104) | 0.955 (0.05) | 0.623 (0.117) |
|  |  | Iberia | 0.851 (0.029) | 0.710 (0.081) | 0.925 (0.018) | 0.506 (0.089) |
|  |  | Italy | 0.794 (0.036) | 0.760 (0.072) | 0.884 (0.016) | 0.497 (0.140) |
|  |  | Balkans | 0.720 (0.054) | 0.725 (0.065) | 0.877 (0.019) | 0.418 (0.110) |
|  | Eastern Europe | — | 0.840 (0.023) | 0.653 (0.046) | 0.897 (0.018) | 0.497 (0.060) |
| South Asia | — | — | 0.988 (0.004) | 0.964 (0.009) | 0.992 (0.004) | 0.946 (0.009) |
| Sub-Saharan Africa | — | — | 0.982 (0.005) | 0.993 (0.004) | 0.988 (0.004) | 0.990 (0.002) |
|  | Central-South Africa | — | 0.973 (0.009) | 0.928 (0.044) | 0.997 (0.004) | 0.890 (0.057) |
|  | West Africa | — | 0.964 (0.007) | 0.978 (0.004) | 0.974 (0.005) | 0.962 (0.008) |
|  | East Africa | — | 0.919 (0.016) | 0.938 (0.014) | 0.950 (0.011) | 0.889 (0.022) |
| Middle East and Northern Africa | — | — | 0.940 (0.008) | 0.826 (0.051) | 0.949 (0.008) | 0.764 (0.080) |
|  | Northern Africa | — | 0.950 (0.011) | 0.772 (0.045) | 0.953 (0.011) | 0.711 (0.081) |
|  | Middle East | — | 0.846 (0.017) | 0.865 (0.040) | 0.895 (0.014) | 0.764 (0.080) |

At the regional scale, precision, recall and coverage are uniformly equal to or lower than in the continental-scale parent populations, by definition (Table 1). With *t* = 0, precision remains fairly high, above 84% for all regions, and over 90% for most. Recall for *t* = 0 is relatively good at this scale, well above 90% for most regions, but falls off for several regions, including Eastern Europe.

When the precision threshold is increased to *t* = 0.8, average precision climbs to 90% for all regions, and to 95% or above for most. Recall for *t* = 0.8 falls for all populations, ranging from about 65% to 95% across regions, with the exception of Eastern Europe, where it falls to just under 50%. The combination of high precision and moderately high recall suggests that the consequence of the increased precision threshold is modest improvement in the true and false positive rates, with larger increase in the false negative rate. The false negative rate can climb across all the regions at once, because segments that go unassigned at the *t* = 0.8 threshold are false negatives for every population; being a false negative with respect to one population does not entail being a false positive with respect to another.

At the sub-regional scale, some populations, such as China, Japan, and Finland, continue to have high precision and recall, for both precision thresholds (Table 1). Again, precision and recall generally decline relative to the regional parent groups. The effect of the precision threshold is much greater at the sub-regional scale that at the two larger scales. Increasing the threshold *t* from 0 to 0.8 typically increases the average precision by 8-20% in populations that do not already show high precision at *t* = 0. The attendant drop in recall ranges from about 4 to 40%.

Within Europe, the France/Germany population has unusually low recall, at about 8%, while populations around Europe’s “periphery” exhibit higher recall. The precision and recall for the Northern Europe subregional populations appears to be lower in general than for the Southern Europe sub-regional populations.

Precision for “France/Germany” is reported at 78.1%, which would appear to be inconsistent with the application of an *t* = 0.8 precision threshold. The recalibration module ensures that the posterior probabilities are well-calibrated at any precision threshold for each population, on average. In this case, it may be that it is relatively uncommon to encounter posterior probabilities much in excess of 80% for France/Germany, as is consistent with the low recall. That the estimated precision is underneath 80%, then, is likely sampling error.

The above tables illustrate that Ancestry Composition achieves high precision and recall at the continental scale as well as the regional scale. At the sub-regional scale, several populations achieve precision and recall greater than 90%. The contrast between the two precision threshold choices shows that due to the recalibration step, Ancestry Composition can be tuned to achieve a desired precision, at the cost of recall and coverage. For example, the low coverage estimate for Scandinavia means that if one wanted to achieve 80% precision, one would typically assign 34.5% of a genuinely Scandinavian individual’s genome to Scandinavia.

### 3.3 Ancestry Composition validation

In order to further validate Ancestry Composition, we estimated the genome-wide proportions of African, European and Native American ancestry in admixed populations from the 1000 Genomes project. In particular, we ran Ancestry Composition on 61 African-American individuals (ASW population), 60 Colombians (CLM), 66 Mexican (MXL) and 55 Puerto-Rican individuals (PUR). These individuals’ global admixture proportions were previously estimated using a consensus of four published admixture deconvolution methods (The 1000 Genomes Project Consortium, 2012).

We detected four ASW individuals with high proportions of Native American ancestry (NA20299, NA20314, NA20414 and NA19625). The first three of these individuals have previously been reported as likely to have Native American ancestry (Gourraud *et al.*, 2014). Ancestry Composition was run over the test individuals considering all 25 of its reference populations simultaneously, so allowed the possibility of assignment to any of those populations. We note, by contrast, that the 1000 Genomes results for the ASW population were obtained using two-way (European and African) admixture estimates. Therefore, the Native American component of ASW ancestry is always zero by definition. As a consequence, the 1000 Genomes consensus local admixture inference forces segments of Native American ancestry to be of either European or African origin; our results suggest that such segments are mostly called as European in the consensus estimates, as opposed to African. Table S4 reports the mean African, European and Native American ancestries estimated with Ancestry Composition and the consensus 1000 Genomes analysis in each of the four populations considered. To assess the agreement between the Ancestry Composition and 1000 Genomes estimates, we fit a linear regression model

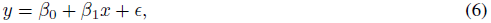

where *y* are the admixture proportions estimated by the 1000 Genomes consensus method, *x* are the admixture proportions estimated by Ancestry Composition, and *β*_0_*, β*_1_ are the coefficients of the regression. Table S4 reports the estimated values of *β*_0_ and *β*_1_ as well as the coefficient of determination *r*^2^. In every case, the agreement between Ancestry Composition estimates and the 1000 Genomes consensus method estimates is very high, with *r*^2^ values higher than 0.9 and *β*_1_’s very close to 1. Figure S4 plots the Ancestry Composition estimates as a function of the 1000 Genomes consensus estimates. We note that all points fall on the first diagonal at a default precision threshold *t* = 0.90, with the exception of Native American estimates in the CLM, MXL and PUR populations. In this case, Ancestry Composition estimates lower amount of Native American ancestry than the 1000 Genomes consensus method. We note that lowering the precision threshold to *t* = 0.51 recovers the 1000 Genomes consensus method estimates. Thus, Ancestry Composition appears to be conservative in its estimates of Native American ancestry, possibly due to the relatively small reference population used.

## 4 Discussion

We have developed a new, modular pipeline for ancestry deconvolution. The pipeline is composed of three stages that were developed as independent modules. We believe the modular approach we have taken makes Ancestry Composition flexible, robust, and easy-to-update. Our cross-validation experiments showed that Ancestry Composition achieves high precision and recall for populations separated by continental and subcontinental-scale distances. Unlike previous approaches, generally restricted to a few well-differentiated populations, Ancestry Composition is able to handle a large number of closely-related populations simultaneously.

### Limits of assignment resolution

Our results indicate near-perfect precision and recall at the continental scale, with declining precision and recall at the regional and sub-regional scales. In Europe, the classifier is usually able to distinguish Northern from Southern from Eastern European haplotypes, but encounters difficulty at the sub-regional, let alone the national level. At the sub-regional level, the classifier exhibits poor coverage in the “France/Germany” reference population that includes samples from many central European nations. The coverage is higher in populations away from Europe’s center, such as Britain/Ireland, Scandinavia, and Iberia, and at similar precision levels.

There are two logical possibilities when the classifier fails to assign a haplotype to its correct reference population. The first is that the assignment should have been made, but the classifier as implemented made a mistake due to insufficient data, random sampling, or a failure of one or more of its modeling assumptions. The second, perhaps more intriguing, possibility is that the assignment is not there to be made.

This pattern in our coverage estimates might reflect the fact that some haplotypes are relatively private to a geographic region, and some haplotypes are more cosmopolitan. Culturally- and/or geographically-isolated populations like Finns and Ashkenazi Jews may exhibit high coverage due to a greater proportion of relatively private haplotypes, while highly-connected central European populations may have relatively few private haplotypes.

Therefore, while it is likely that precision, recall and coverage would increase with a larger, more robust dataset and with refinements to our model, it is possible that there are limits to assignment coverage because some proportion of haplotypes are fundamentally shared between populations.

### Discriminative versus generative approaches

The goal of ancestry deconvolution is ultimately to estimate *P* (*Y* | *X*), the distribution of the unobserved ancestry states *Y* given the observed haplotypes *X*. Generative approaches, such as HMMs, first estimate the joint distribution *P* (*X, Y*) before conditioning on the observation *X*. By contrast, discriminative approaches directly model the conditional distribution *P* (*Y* | *X*). Discriminative approaches are well suited for classification where they can outperform generative methods (Lafferty *et al.*, 2001). The additional complexity required to model *P* (*X, Y*) fully also limits most generative ancestry deconvolution methods to a few ancestral populations.

By not attempting to model the joint distribution of haplotypes and their ancestry fully, discriminative approach are typically more robust to model misspecifications. Indeed, our support vector machine local classifier does not assume a particular demographic model underlying the admixture process; instead, it attempts to learn boundaries between the different reference populations directly from the data.

Ancestry Composition adopts a mixed approach, in which the output of a discriminative local classifier is input into a generative error correction module implemented as an autoregressive pair hidden Markov model. Generative models are generally more flexible in expressing complex dependencies between the observation and the hidden random variables. Indeed, the hidden Markov framework provided a natural means to correct for switch errors and allowed a natural extension to model the dependencies between adjacent observations. We note that the idea of combining discriminative and generative models has been employed elsewhere (Jaakkola *et al.*, 1999; Ng and Jordan, 2002).

A purely discriminative approach from admixture deconvolution was published recently (Maples *et al.*, 2013). It implements random forests as its local classifier and conditional random fields to reconcile adjacent chromosomal segments. Random forests have an advantage over SVMs that they are multi-class classifiers and their output is fundamentally probabilistic in nature. However, SVMs offer a direct way to plug in kernels, which enabled us to extract a huge number of features from short chromosomal segments without compromising the speed of Ancestry Composition.

### Classification with reliable probabilities

We note that Ancestry Composition is the first local admixture inference method to use a recalibration method. Recalibrating Ancestry Composition’s posterior probabilities has at least two advantages. First, it enables us to easily tune the recall/precision tradeoff. Indeed, once the posterior probabilites have been recalibrated, they directly translate into the precision achieved by the classification. Second, and perhaps more importantly, recalibration is a natural framework for reconciling classification of haplotypes with the fact that many haplotypes are not specific to a population (Novembre and Ramachandran, 2011). As an illustrative example, let us assume that a haplotype has posterior probabilities of 34%, 33% and 33% of belonging to three distinct European sub-populations, A, B and C. Assume we set the desired level of precision to 0.9. Non-recalibrated methods would either assign the haplotype to sub-population A or decide the segment is unassigned. Instead, recalibration enables us to assign to a “higher-level” population, such as “Northern Europe” or “Europe", and guarantees that the precision of the higher-level assignment meets the threshold.

### Reliable admixture deconvolution with many closely related populations

In this work, we demonstrated that the combination of large reference populations and modern machine learning techniques allows to accurately infer local admixture from many reference populations on a subcontinental scale. In addition, we showed the importance of obtaining reliable assignment probabilities, which can then be directly used to tune the precision/recall compromise. To the best of our knowledge, we are the first to use probability calibration techniques in the context of local admixture inference.

## Acknowledgments

We would like to thank Brian Naughton, Katarzyna Bryc, and Nicholas Eriksson for insightful comments throughout this project. We also thank the many 23andMe customers who answered survey questions and who allowed us to study their genomes, without whom this work would not have been possible.

## List of Figures

1 **Overview of Ancestry Composition pipeline.** An individual genotype is first phased using an out-of-sample extension of BEAGLE, which outputs two haplotypes, *hap*_1_*, hap*_2_. The haplotypes are passed to the local classifier, where they are split in equal size chunks that are classified independently to one of the reference populations. The local classifier outputs two hard-clustering vectors, *hc*_1_*, hc*_2_, composed of populations of origin on each haplotype. The error correction module takes *hc*_1_*, hc*_2_ as input, reconciles hard-clustering calls and corrects for switch errors. It outputs two vectors, *p*_1_*, p*_2_, containing assignment probabilities along both haplotypes. Finally, *p*_1_ and *p*_2_ are fed to a recalibration module that outputs assignment.

## List of Tables

1 **Ancestry Composition accuracy at all scales.** Precision and recall are shown for the 31 populations, evaluated at precision thresholds of *t* = 0 and *t* = 0.8. A threshold of *t* = 0 corresponds to making best-guess predictions. The values in parentheses are the respective standard errors computed across chromosomes. The “—” symbol indicates that precision and recall values are integrated over all children of the parent population.

